# Transposable elements and gene expression during the evolution of amniotes

**DOI:** 10.1101/283390

**Authors:** Lu Zeng, Stephen M. Pederson, R. Daniel Kortschak, David L. Adelson

## Abstract

**Background:** Transposable elements (TEs) are primarily responsible for the changes in genome sequences that occur over time within and between species. TEs themselves evolve, with clade specific LTR/ERV, LINEs and SINEs responsible for the bulk of species specific genomic features. Because TEs can contain regulatory motifs, they can be exapted as regulators of gene expression. While TE insertions can provide evolutionary novelty for the regulation of gene expression, their overall impact on the evolution of gene expression is unclear. Previous investigators have shown that tissue specific gene expression in amniotes is more similar across species than within species, supporting the existence of conserved developmental gene regulation. In order to understand how species specific TE insertions might affect the evolution/conservation of gene expression, we have looked at the association of gene expression in six tissues with TE insertions in six representative amniote genomes (human, opossum, platypus, anole lizard, bearded dragon and chicken).

**Results:** We have used a novel bootstrapping approach to minimise the conflation of effects of repeat types on gene expression. We compared the expression of orthologs containing different types of recent TE insertions to orthologs that contained older TE insertions and found significant differences in gene expression associated with TE insertions. Likewise, we compared the expression of non-ortholog genes containing different types of recent TE insertions to non-orthologs with older TE insertions and found significant differences in gene expression associated with TE insertions. As expected TEs were associated with species-specific changes in gene expression, but the magnitude and direction of change of expression changes were unexpected. Overall, orthologs containing clade specific TEs were associated with lower gene expression, while in non-orthologs, non clade-specific TEs were associated with higher gene expression. Exceptions were SINE elements in human and chicken, which had an opposite association with gene expression compared to other species.

**Conclusions:** Our observed species-specific associations of TEs with gene expression support a role for TEs in speciation/response to selection by species. TEs do not exhibit consistent associations with gene expression and observed associations can vary depending on the age of TE insertions. Based on these observations, it would be prudent to refrain from extrapolating these and previously reported associations to distantly related species.

## Introduction

Transposable Elements (TEs) have been shown to alter gene regulation and drive genome evolution [1] [2] [3] [4]. TEs can exert these effects on genes by altering chromatin structure, providing novel promoters or insulators, novel splice sites or other post-transcriptional modifications to re-wire transcriptional networks important in development and reproduction [2] [5]. TEs that land in introns can become “exonized” or spliced into mRNA of the gene into which they have inserted, often introducing stop codons into mRNA that can lead to nonsense-mediated mRNA decay, serving to control gene expression [6] [7].

Short INterspersed Elements (SINEs) are non-autonomous TEs ancestrally related to functionally important RNAs, such as tRNA, 5S rRNA and 7SL RNA that replicate by retrotransposition. SINEs possess an internal promoter that can be recognized and transcribed by the RNA polymerase III (polIII) enzyme complex, and are usually present in a monomeric or tandem dimeric structure [8]. Monomeric tRNA-related SINE families are present in the genomes of species from all major eukaryotic lineages and this structure is, by far, the most frequent. These elements are composed of a 5’ tRNA-related region and a central region of unknown origin, followed by a stretch of homopolymeric adenosine residues or other simple repeats [9] [10]. In contrast to the very widespread phylogenetic distribution of tRNA derived SINEs, 7SL-derived SINEs have been found only in mammals [8]. They are composed of a 7SL-derived region followed by a poly(A) tail and can be either monomeric (B1 family) or dimeric (Alu family) [11] [12]. 5S rRNA-derived SINEs were found in fishes (SINE3) but were likely active in the common ancestor of vertebrates [13] [14]. They are with a 5S-related region (instead of a tRNA-related region), followed by a central region of unknown origin and 3-terminal repeats [13]. SINE RNAs have also been shown to possess the potential to regulate gene expression at the post-transcriptional level, for example, Alu RNAs an modulate protein translation, influence on RNA editing and mRNA splicing [15].

Long INterspersed Elements (LINEs) are autonomously replicating TEs that replicate through an RNA intermediate that is reverse transcribed back into the genome at a new location. LINEs contain an internal DNA Polymerase II promoter and either one or two Open Reading Frames (ORFs) that contain a Reverse Transcriptase (RT) domain and an Endonuclease (EN) domain. L1 family repeats show a stronger negative correlation with expression levels than the gene length [16], and the presence of L1 sequences within genes can lower transcriptional activity [17].

Long terminal repeat (LTR) retrotransposons are a group of TE, that are flanked by long terminal repeats and contain two ORFs: *gag* and *pol*. The *gag* ORF encodes the structural protein that makes up a virus-like particle [18]. The pol ORF encodes an enzyme needed for replication that contains protease, integrase, reverse transcriptase, and RNase H domains required for reverse transcription and integration. LTRs can also act as alternative promoters to provide new tissue-specificity, act as the major promoters, or exert only minor effects [19]. Many endogenous retroviruses (ERV) contain sequences that can serve as transcriptional start sites or as cis-acting regulatory elements in the host genomes [20].

DNA transposons encode a transposase gene that is flanked by two Terminal Inverted Repeats (TIRs) [21]. The transposase recognizes these TIRs to excise the transposon DNA, which is then inserted into a new genomic location by cut and paste mobilization[22]. DNA transposons can inactivate or alter the expression of genes by insertion within introns, exons or regulatory region [1] [21].

There is a growing realization that many TEs are highly conserved among distantly related taxonomic groups, suggesting their biological value to the genome. In this report, we describe the association of clade specific TEs with gene expression in long diverged amniotes (Figure 1A) in order to determine how much these TEs might have altered the regulation of gene expression in six tissues during the evolution of these species.

**Figure 1:**
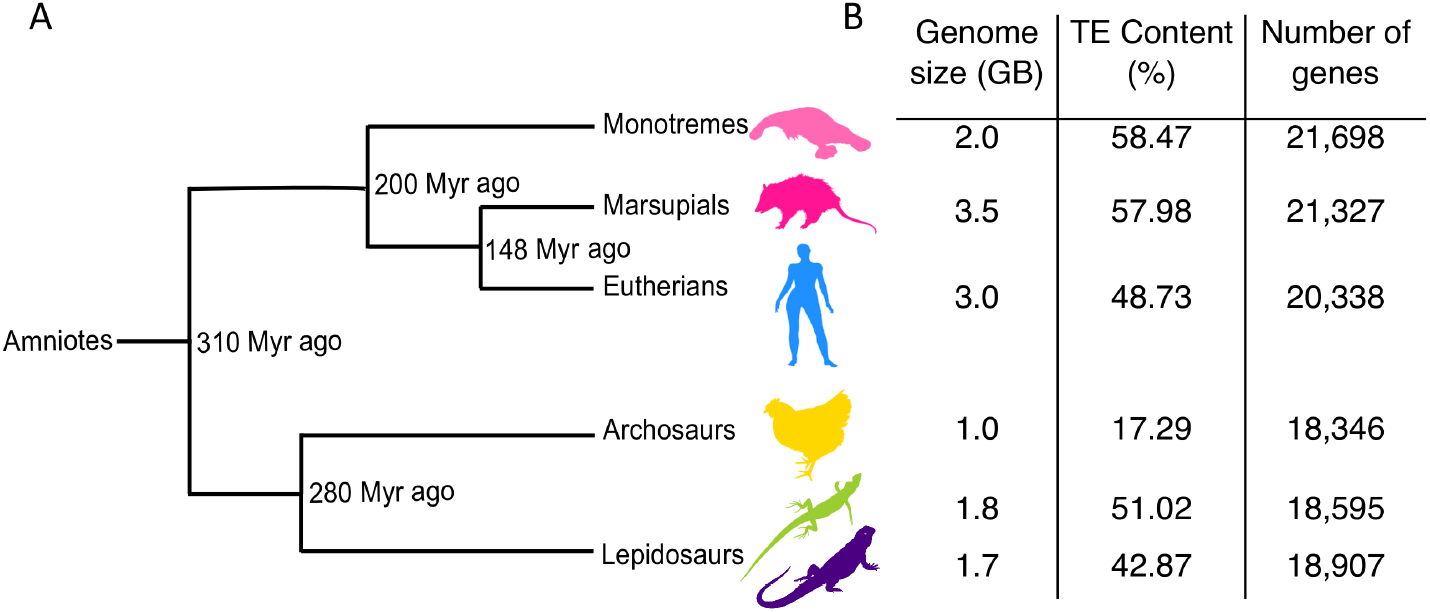
Divergence times and genome statistics of the major amniote lineages. A) The silhouettes indicate species used in this study: *Ornithorhynchus anatinus* (platypus), *Monodelphis domestica* (opossum), *Homo sapiens* (human), *Gallus gallus* (chicken), *Anolis carolinensis* (anole lizard) and *Pogona vitticeps* (bearded dragon) (from top to bottom). Time since main speciation events obtained from TimeTree (www.timetree.org) are indicated (millions of years ago, Myr ago) [24]; B) Genome statistics.

## Methods

### Expression data

RNA-seq expression data were available for six species (Table 1), belonging to the five main amniote lineages (eutherian: human; marsupial: gray short-tailed opossum; monotreme: platypus; lepidosaur: green anole lizard, bearded dragon; archosaur: chicken) from four somatic (brain, heart, liver, kidney) and two reproductive tissues (testis, ovary)(Gene Expression Omnibus accession numbers GSE30352 [23] and GSE97367 [24], BioProject number PRJEB5206 [25]).

**Table 1:** Summary of datasets and tissue samples analyzed in this study.

| Dataset(s) | Tissues | Species |
| --- | --- | --- |
| Marin | brain, heart, kidney, liver, ovary, testes | chicken, anole, platypus, opossum, *human |
| Brawand | brain, heart, kidney, liver, testes | chicken, anole, platypus, opossum, human |
| Private | brain, heart, kidney, liver, ovary, testes | bearded dragon |

\* Human samples in this set do not include ovary tissue.

Trim_galore (v0.4.2)(–clip_R1 5; –three_prime_clip_R1 5) [26] was used for adapter trimming and quality control. Adapter-trimmed RNA-seq reads were aligned to the reference genomes (Ensembl release 74) with RSEM (v1.3.0) [27] using Bowtie2 (v2.2.9) [28] with default parameters as the alignment tool. Gene expression was estimated as TPM (Transcripts Per Million). A complete list of accessions can be found in Table S1, Additional file 1.

### Genomic data

For chicken, anole lizard, platypus, opossum and human, gene annotations were download from Ensembl release 74. For bearded dragon, RefSeq assembly GCF_900067755.1 was used for analysis. Complete information on genomes used can be found in Table S2, Additional file 1.

### Ortholog definition

Gene orthologies were downloaded from Ensembl release 74. Amniote orthologs were defined as single-copy orthologous genes conserved in all 6 amniote species. Reciprocal best hits were used to extract orthologous genes between bearded dragon and other five species by using BLASTN [29]. A total number of 6,595 orthologous genes were extracted from the six species.

### TE annotation

TEs were annotated by using CARP: a *ab initio* method [30]. Recently inserted, low divergence TE referred to hereafter as species-specific TE (ssTE) were defined as having ≥94% sequence identity. They were extracted from CARP output, which identifies and annotates TEs that have ≥94% sequence identity. Older TEs were defined as the remaining TE insertions in the genome and are referred to as non-species specific TE (nsTE).

### The weighted bootstrap procedure for assessing association of gene expression and TEs

Many genes contain multiple transposable elements, with only a minority of genes containing a single TE. In order to assess any effects on transcription due to the presence of a single TE, a weighted bootstrap approach was devised. For a given TE type within each individual gene, the frequencies of co-occurring TE types and combinations of TE types were noted. Uniform sampling probabilities were then used for the set of genes containing a specific TE type (test sample), whilst sampling weights were assigned to genes lacking the specific TE type based on TE composition (reference sample) (See detail in Table S3-6, Additional file 1). Gene length was divided into 10 bins and these were included as an additional category when defining sampling weights. This ensured that two gene sets were obtained for each bootstrap iteration, which were matched in length and TE composition with the sole difference being the presence of the specific TE type. The median difference in expression level, as measured by log2(TPM), and the difference in the proportions of genes detected as expressed were then used as the variables of interest in the bootstrap procedure. The bootstrap was performed on sets of 1,000 genes (except for ortholog genes containing non-recent species specific SINE elements in platypus) for 5,000 iterations. Samples that could not meet the minimum number of 600 genes were not used. When comparing expression levels, genes with zero read counts were omitted prior to bootstrapping. In order to compensate for multiple testing considerations, confidence intervals were obtained across the m=nTissues*nElements tests at the level 1-/m, which is equivalent to the Bonferroni correction, giving confidence intervals that controlled the family-wise error rate at the level =0.05. Approximate two-sided p-values were also calculated by finding the point at which each confidence interval crossed zero, and additional significance was determined by estimating the FDR on these sets of p-values using the Benjamini-Hochberg method.

## Results

### Mammalian gene expression phylogenies

To obtain an initial overview of gene expression patterns, we evaluated the similarity of ortholog gene expression in 6 tissues (heart, brain, kidney, liver, testis and ovary), from both males and females in our 6 species. These RNA-seq samples were assembled from three different studies (Table 1, further detail can be seen in Additional file Table S1) [24] [23] [25].

Two hierarchical clustering methods were used to investigate the conservation of expression signatures in these six species within six tissues. 1 - Unweighted Pair Group Method with Arithmetic Mean (UPGMA) and 2 - Ward’s minimum variance (ward.D2) hierarchical clustering.

While mostly similar, the two methods did give slightly different clustering results (Figure 2). Generally, gene expression clustered according to tissue with three exceptions. The first exception was bearded dragon heart expression clustered using Ward’s method, where heart samples clustered with kidney and liver samples. The second exception was for platypus testis expression clustered using UPGMA, where testis expression clustered with ovary. The third exception was more widespread, and found with both clustering methods; kidney and liver samples only clustered by tissue for human and opossum and were found together more often in species-specific clusters for the other species.

**Figure 2:**
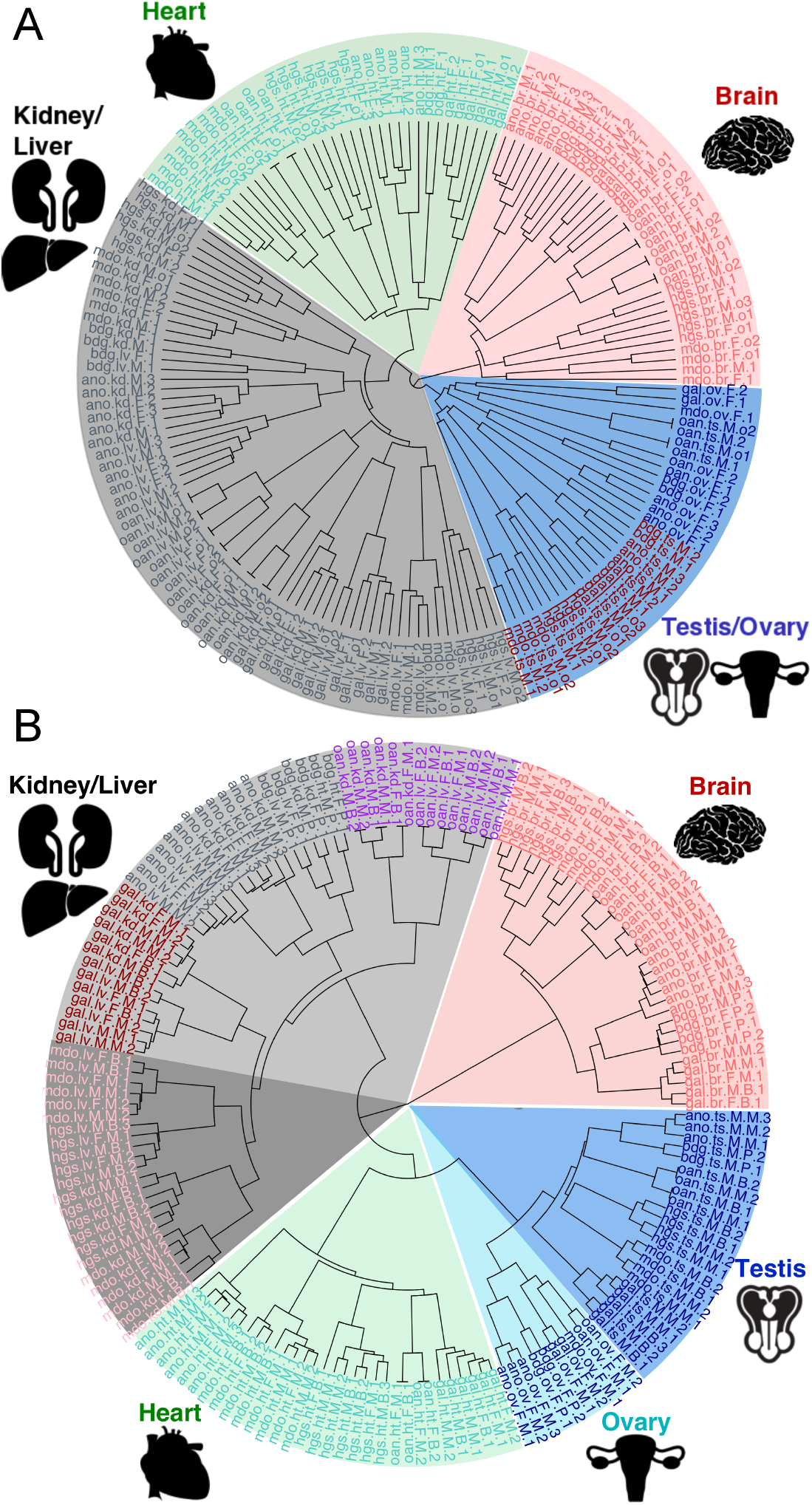
Tissue specific *vs* species specific clustering of gene expression in amniotes. a, Clustering of samples based on expression values, calculated as transcripts per million (TPM) of one to one orthologous genes expressed in heart, brain, kidney, liver, testis and ovary (n=6596). UPGMA (Unweighted Pair Group Method with Arithmetic Mean) hierarchical clustering was used with distance between samples calculated using the average of all distances between pairs. b, Clustering of samples based on expression values, calculated as transcripts per million (TPM) of one to one orthologous genes expressed in heart, brain, kidney, liver, testis and ovary (n=6596). Ward’s minimum variance hierarchical clustering was used with distance between samples measured by the squared Euclidean distance.

### Comparison of gene expression for genes on the basis of their TE content

There were two aspects of the data that affected our analysis. First, because the vast majority of genes contain TEs, it was impossible to compare expression of genes with TEs against genes without TEs, as there were too few of the latter. So we designed our comparisons as shown in Figure 3. Second, most genes contain multiple TE types. In
order to minimize the conflation of co-occuring TEs, a weighted bootstrap approach was used in this study. The idea is simple, if we want to investigate the association between a SINE insertion and gene expression, first we randomly select 1,000 genes that contain a SINE element, and then compare their expression level to 1,000 randomly selected genes that do not contain any SINEs. We repeat this process 5,000 times in order to generate enough observations for statistical analysis.

**Figure 3:**
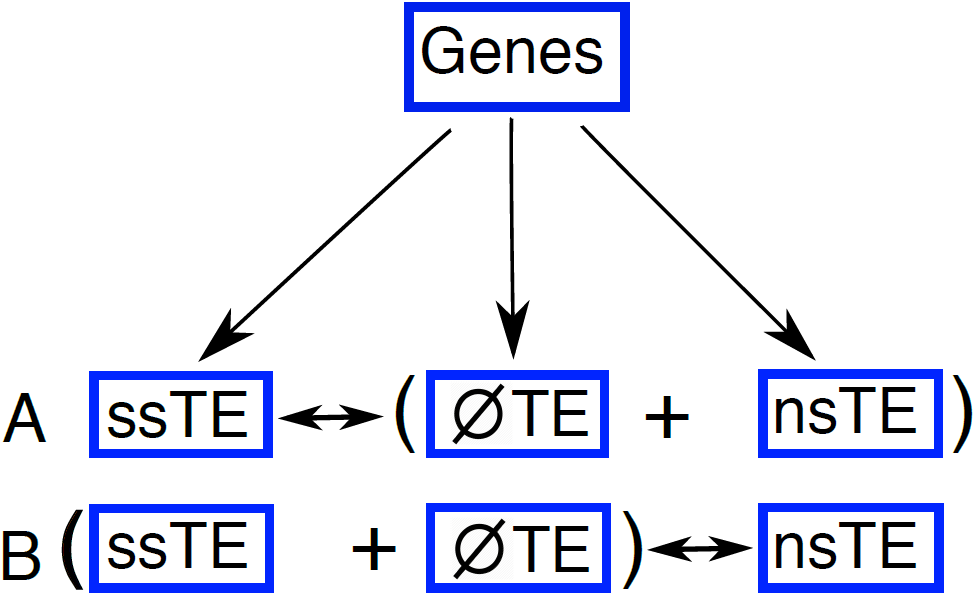
Gene sets for expression comparison. Because there were too few genes (orthologs or non-orthologs) with no TE insertions, we designed comparison sets based on the following scheme. We split our gene sets (either ortholog or non-ortholog) into three subsets: those containing recent species specific TE insertions (ssTE), those containing non-recent species specific TE insertions (nsTE) and those containing no TE (ØTE)

### Ortholog expression is associated with with TE type

For our specific analyses, BedTools was used to get the intersection between TE types and 6,595 orthologous genes (including 1kb upstream and 1kb downstream regions) within our six species (chicken, anole lizard, bearded dragon, platypus, opossum and human). The boostrap approach as described above was then applied to this data in order to investigate the association between orthologous gene expression and TE insertions. TEs were split into two groups: recently inserted, low divergence TEs, referred to as species-specific TEs (ssTEs, see methods for detail) and more divergent TEs, referred to as non-species specific TEs (nsTEs). Genes containing no TEs are referred to as ØTE. The two TE groups were further broken down into four TE classes: DNA transposon, ERV/LTR, LINE or SINE.

Because purifying selection is likely to be more common on orthologs, and since tissue specificity of ortholog expression was largely conserved (Figure 2), we looked first at the association ortholog expression with TE insertions. We compared expression for orthologs containing ssTE against orthologs containing nsTE + ØTE and expression of orthologs containing nsTE against orthologs containing ssTE + ØTE (Figure 4) and (Figures S1 and S2, Additional file 1). We found that ssTEs (ERV/LTR, LINE and SINE) were associated with lower gene expression in orthologs, especially in anole lizard, bearded dragon and human. The exceptions to this negative association were in the human and chicken genome, where recent insertions of SINEs were found associated with higher gene expression in testis and brain.

**Figure 4:**
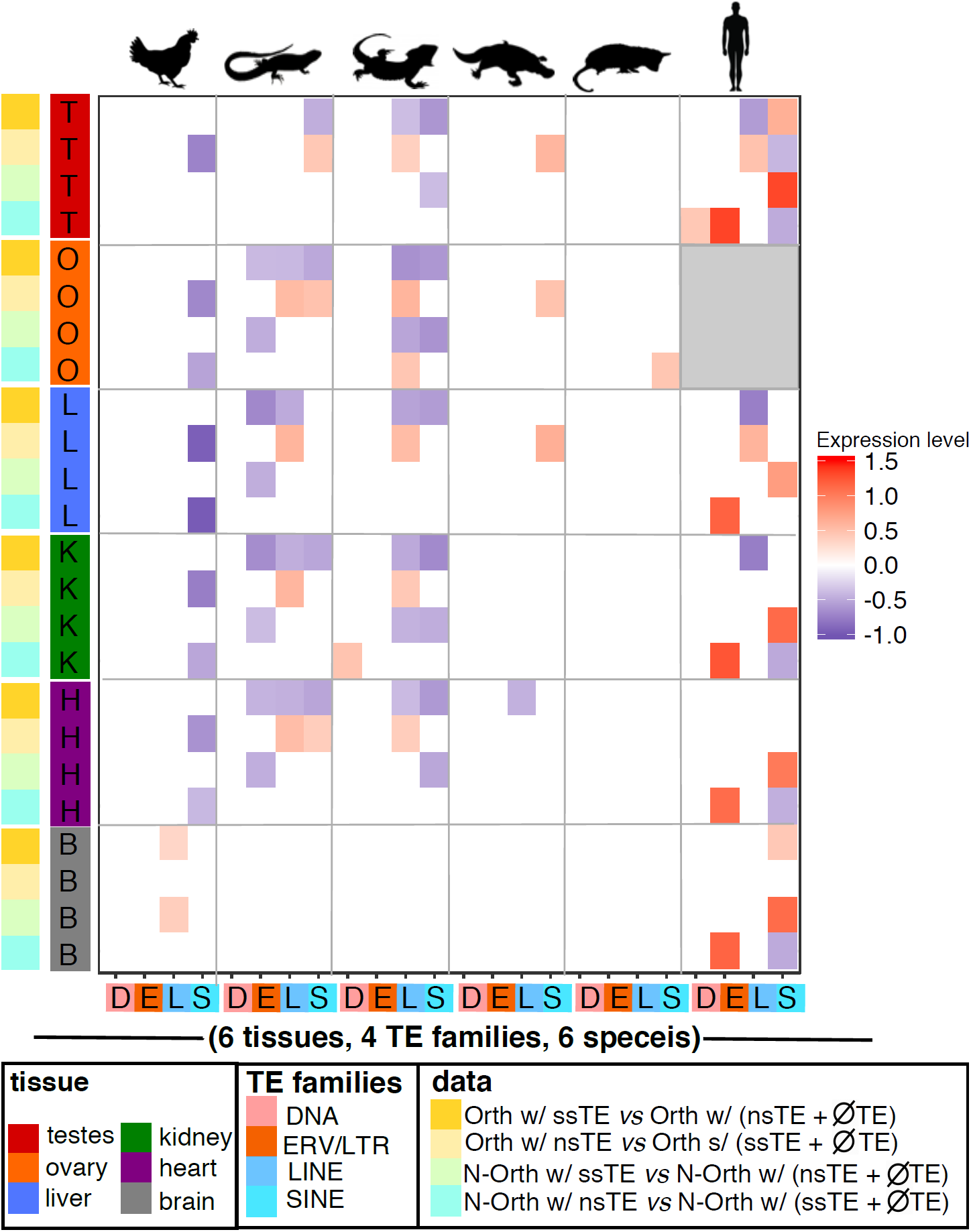
Changes in the levels of ortholog/non-ortholog gene expression as a function of TE insertion. This figure shows the association between orthologous/non-orthologous gene expression levels in six species (from left to right: anole lizard, chicken, human, opossum, platypus and bearded dragon(pogona)) with the presence of recent species-specific TE insertions (ssTE) or non-recent species specific TE insertions (nsTE) (from left to right: LINE, SINE, ERV/LTR or DNA). A weighted bootstrap approach was used to compare the median difference in gene expression levels of orthologous/non-orthologous genes with a ssTE/nsTE insertion compared to orthologous/non-orthologous gene without ssTE/nsTE. Gene expression levels are log2-transformed. Comparisons without statistically significant gene expression changes are shown in white. Statistically significant increased gene expression shown in red and statistically significant decreased gene expression in blue. Grey shading indicates no samples were available for this comparison.

For orthologs containing nsTE (LINE or SINE) (Figure 4, additional figure S1 and S3) we observed primarily a positive association with gene expression in contrast to the trend seen with ssTEs. The exceptions to this positive association were again found in the human and chicken genomes. Particularly in the chicken genome, where the insertion of non-species specific SINEs were associated with lower ortholog gene expression in multiple tissues.

Overall, species specific TE insertions in orthologs were mainly associated with lower gene expression, while non-species specific TE insertions were mainly associated with higher gene expression. This is true for ERV/LTR in anole lizard, bearded dragon and human, LINE and tRNA derived SINE insertions in anole, bearded dragon, platypus and human. There are some exceptions, notably for chicken orthologs with nsTE insertions which showed an association with decreased gene expression. Perhaps the most interesting observation was that the magnitude of the effect on gene expression was quite pronounced, ranging between about −30% to +40% changes in median gene expression values (Table S7, Additional file 1).

### Non-ortholog gene expression is associated with TE type

In order to explore the association of TEs in a more general context, we then expanded our analysis from orthologous genes to non-orthologous genes.

As described above BedTools was used to get the intersection between TE types and non-orthologous genes, and the bootstrap approach was used to compare expression for non-orthologs containing ssTE against non-orthologs containing nsTE + ØTE and expression of non-orthologs containing nsTE against non-orthologs containing ssTE + ØTE (Figure 4) and (Figures S4 and S5, Additional file 1).

Similar to orthologs, ssTE insertions in non-orthologs showed a negative association with gene expression. This can be observed in ERV/LTR, LINE and SINE in anole lizard and bearded dragon. In the chicken, older SINE insertions in non-orthologs were negatively associated with gene expression. In contrast to the anole lizard and bearded dragon, where recent ERV/LTR, LINE and SINE insertions were associated with lower gene expression, human (7SL derived) SINE insertions in non-orthologs were strongly associated with higher gene expression. The magnitude of the association of TEs with gene expression was even more pronounced in these comparisons, ranging from about −40% to +2.8x (Figure 4) and (Figures S4 and S6, Table S7, Additional file 1).

## Discussion

Tissue *vs* species clustering of ortholog gene expression had previously been reported using PCA based analysis and used to support the notion that conservation of developmental gene expression programs results in tissue specific gene expression clustering [23] [31] [32]. These results have been reported for single experiments. We did not see quite as compelling tissue clustering of gene expression using PCA on data from aggregated experiments (Figure S7, Additional file 1). However we did see largely similar results when we applied hierarchical clustering methods across the aggregated data (Figure 2). However, in contrast with previous studies, we found liver and kidney gene expression clustered more by species. We attribute this to species specific metabolic adaptations responding to more pronounced environmental selection. We therefore expected to see species specific TE insertions associated with species specific changes in gene expression. For recent species specfic SINE, ERV/LTR and LINE insertions this is precisely what we found. However, we found no tissue specific patterns of association of gene expression with TEs.

We expected species specific TE insertions to be associated with changed gene expression, as they would both alter the spacing of pre-existing regulatory motifs and potentially contribute new regulatory motifs [5] [33] [34]. Because random changes in complex systems usually break things, we expected recent TE insertions to be associated with lower gene expression. While this expectation was largely met, there were some significant exceptions, such as human SINE, which were associated with increased gene expression (see discussion below). Conversely, it has been shown that older TE insertions contribute to re-wiring of transcriptional networks [35] [36] and thus would have had time to be exapted as enhancers and might be associated with increased gene expression. Previous studies have found that differential decay of ancestral TE sequences across species may result in species-specific transcription factor binding sites [37]. This expectation was also met for human ERV/LTR. However to our surprise, older TE SINE insertions in the chicken were associated with decreased gene expression.

We expected the magnitude of changes in gene expression associated with TE insertions to be modest, however our analysis showed that TE insertions were associated with large changes in gene expression. Based on the median value of changed gene expression from our bootstrap analysis, most statistically significant log2 transformed changes in gene expression associated with TE were smaller than −0.5 and many were greater than 1.0, indicating a range of −40% to +100% change in median gene expression.

Species-specific TE, behaved differently depending on insertion age and species. The most striking example of this was seen in human with recent SINE insertions associated with increased gene expression and older SINE associated with decreased gene expression. This is consistent with observations that Alu elements have been exapted as transcription factor binding sites, and highly and broadly expressed housekeeping genes are enriched for Alus [38] [39] [40]. This was in contrast to an opposite relationship with LINE insertion age and expression change in human, but consistent with previously reported accumulation differences for SINE and LINE insertions in mammalian regulatory regions/open chromatin [41]. Furthermore, LINEs behave similarly in reptiles and human, with new LINEs associated with lower gene expression and older LINEs associated with higher gene expression. This suggests similar constraints on accumulation of TE in lizards and mammals. Finally, TEs had the fewest associations with gene expression in opossum and platypus. This might indicate that these two species are better at repressing TE activity than human, lizards and chicken.

## Conclusions

The large changes in gene expression associated with TEs, and the species specific associations of TEs with gene expression support a role for TEs in speciation/response to selection by species. TE types do not exhibit consistent associations with gene expression and observed associations can vary depending on the age of TE insertions. Based on these observations, it would be prudent to refrain from extrapolating these and previously reported associations to distantly related species.

## Acknowledgements

We would like to thank Terry Bertozzi and Catisha Coburn for taking the time to read the manuscript in full and offer helpful comments. This paper would not be possible without the helpful discussion with Zhipeng Qu, and extraodinary IT support from Matt Westlake.

## Additional Files

### Additional file 1 — Supplementary Information

Additional file contains supplementary figures and tables as referred to in the main body of the paper.

