## Supplementary Materials for "Transposable elements and gene expression during the evolution of amniotes"

### List of Figures

S1 **Change in the levels of ortholog gene expression as a function of TE insertion.** 4
S2 **Change in the levels of ortholog gene expression as a function of species specific TE insertion.** 5
S3 **Change in the levels of ortholog gene expression as a function of non-species specific TE insertion.** 6
S4 **Change in the level of non-ortholog gene expression as a function of TE insertion.** 7
S5 **Change in the level of non-ortholog gene expression as a function of species specific TE insertion.** 8
S6 **Change in the level of non-ortholog gene expression as a function of non-species specific TE insertion.** 9
S7 **Factorial map of the principal-component analysis of messenger RNA expression levels.** 10

### Supplementary Figures

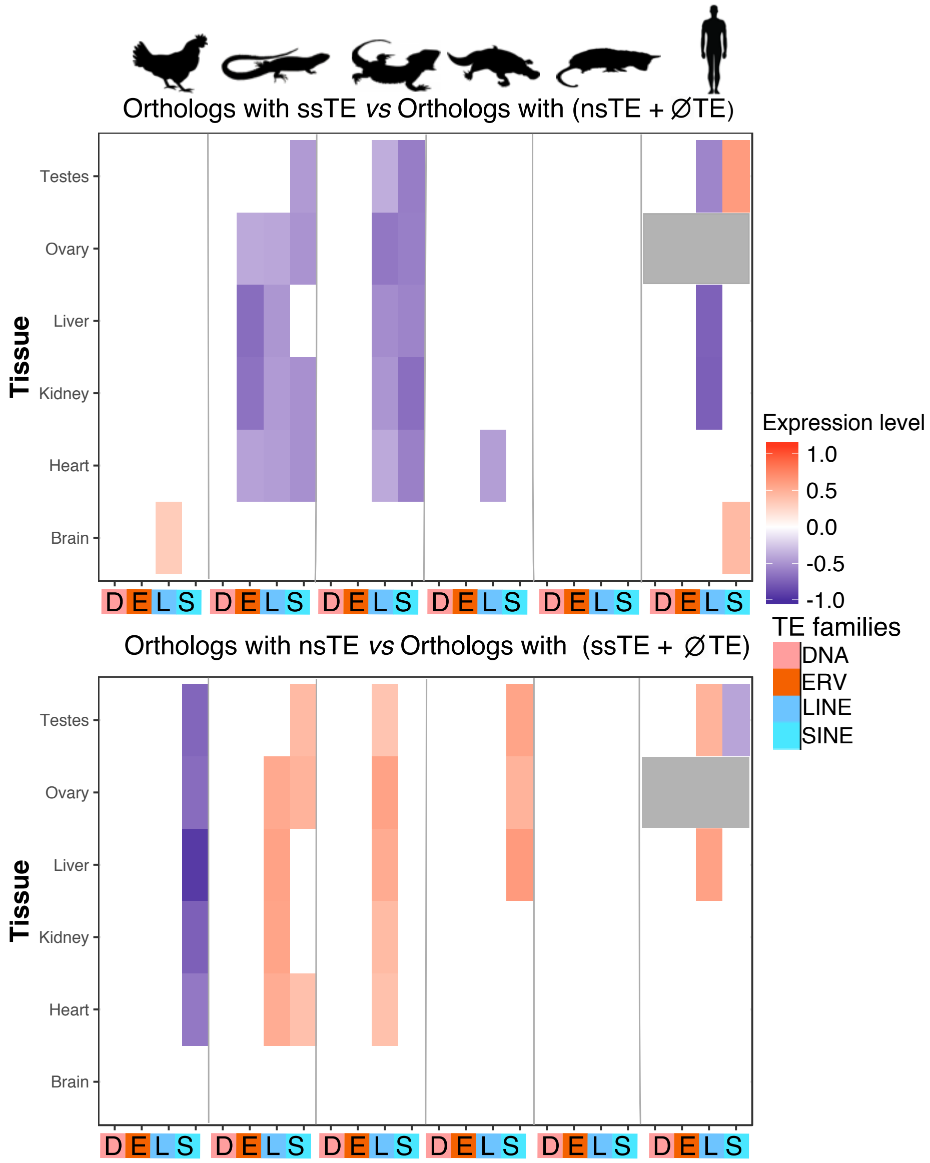

Figure S1: **Change in the levels of ortholog gene expression as a function of TE insertion.** This figure shows the association between ortholog gene expression levels in six species (from left to right: anolis, chicken, human, opossum, platypus and bearded dragon (pogona)) with recent species-specific TE insertions (ssTE) or non-recent species specific TE insertions (nsTE) (from left to right: LINE, SINE, ERV/LTR or DNA). A weighted bootstrap approach was used to compare the median gene expression levels of orthologs with a ssTE/nsTE insertion compared to orthologs without ssTE/nsTE. Gene expression levels are log2-transformed. Comparisons without statistically significant gene expression changes are shown in white. Statistically significant increased gene expression shown in red and statistically significant decreased gene expression in blue. Grey shading indicates no samples were available for this comparison.

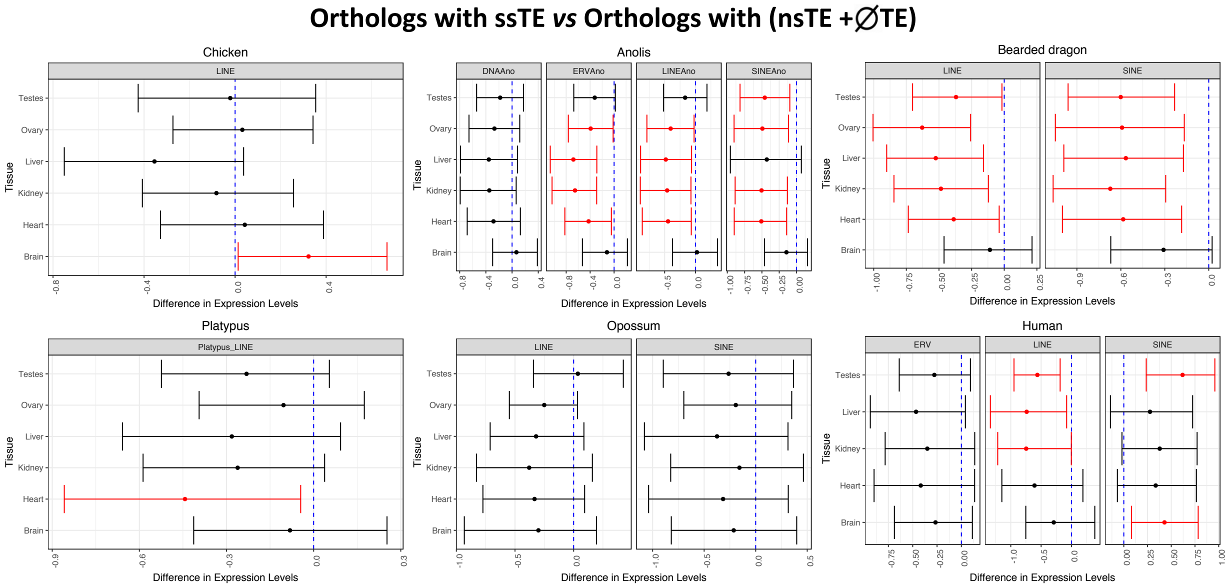

Figure S2: **Change in the levels of ortholog gene expression as a function of species specific TE insertion.**

This figure shows the association between ortholog median gene expression levels in six species (from left to right: anolis, chicken, human, opossum, platypus and bearded dragon (pogona)) with recent species-specific TE insertions (ssTE) (from left to right: LINE, SINE, ERV/LTR or DNA). Confidence Intervals for the difference in median log2(TPM) counts. Confidence Intervals were obtained using the weighted bootstrap and are 1-$\alpha$/m intervals, where $\alpha$=0.05 and m=nTissues*nElements as the total number of intervals presented. Red dots represent the median value from the bootstrap procedure, whilst the vertical line indicates zero. Intervals which do not contain zero are coloured red, and indicate a rejection of the null hypothesis, $H_0$: $\Delta\theta$=0, where $\theta$ represents the parameter of interest.

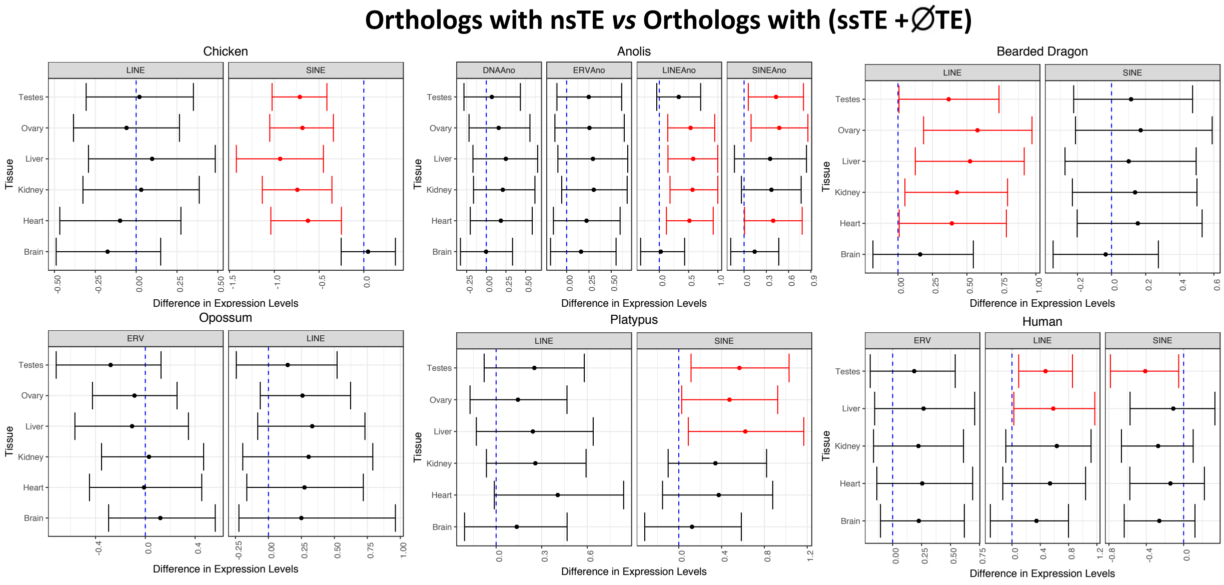

Figure S3: **Change in the levels of ortholog gene expression as a function of non-species specific TE insertion.**
This figure shows the association between ortholog gene expression levels in six species (from left to right: anolis, chicken, human, opossum, platypus and bearded dragon (pogona)) with non-recent species-specific TE insertions (nsTE) (from left to right: LINE, SINE, ERV/LTR or DNA). Confidence Intervals for the difference in median log2(TPM) counts. Confidence Intervals were obtained using the weighted bootstrap and are 1-$\alpha$/m intervals, where $\alpha$=0.05 and m=nTissues*nElements as the total number of intervals presented. Red dots represent the median value from the bootstrap procedure, whilst the vertical line indicates zero. Intervals which do not contain zero are coloured red, and indicate a rejection of the null hypothesis, $H_0$: $\Delta\theta$=0, where $\theta$ represents the parameter of interest.

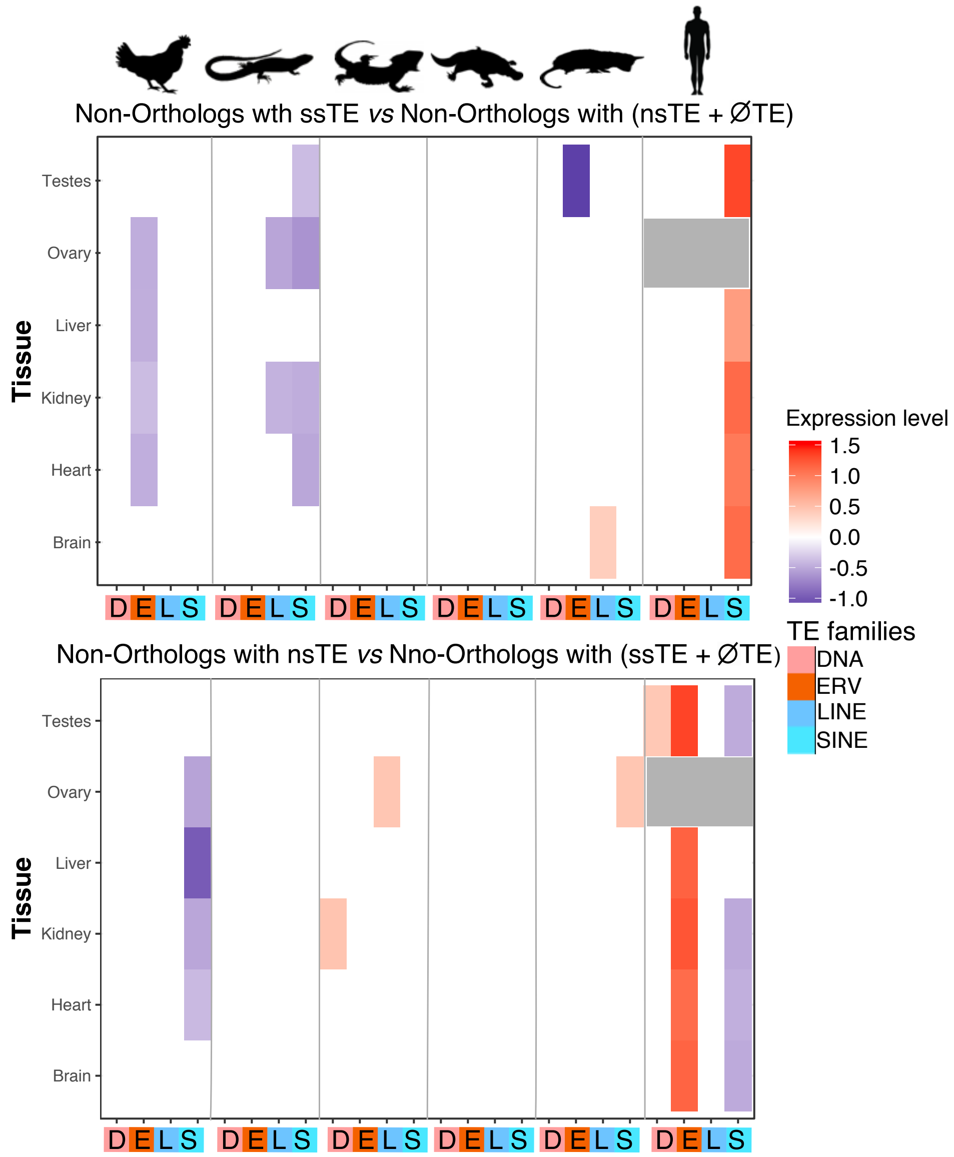

Figure S4: **Change in the level of non-ortholog gene expression as a function of TE insertion.** This figure shows the association between non-ortholog gene expression levels in six species (from left to right: anolis, chicken, human, opossum, platypus and bearded dragon (pogona)) with recent species-specific TE insertions (ssTE) or non-recent species specific TE insertions (nsTE) (from left to right: LINE, SINE, ERV/LTR or DNA). A weighted bootstrap approach was used to compare the median gene expression levels of non-orthologous genes with a ssTE/nsTE insertion compared to non-orthologous gene without ssTE/nsTE. Gene expression levels are log2-transformed. Comparisons without statistically significant gene expression changes are shown in white. Statistically significant increased gene expression shown in red and statistically significant decreased gene expression in blue. Grey shading indicates no samples were available for this comparison.

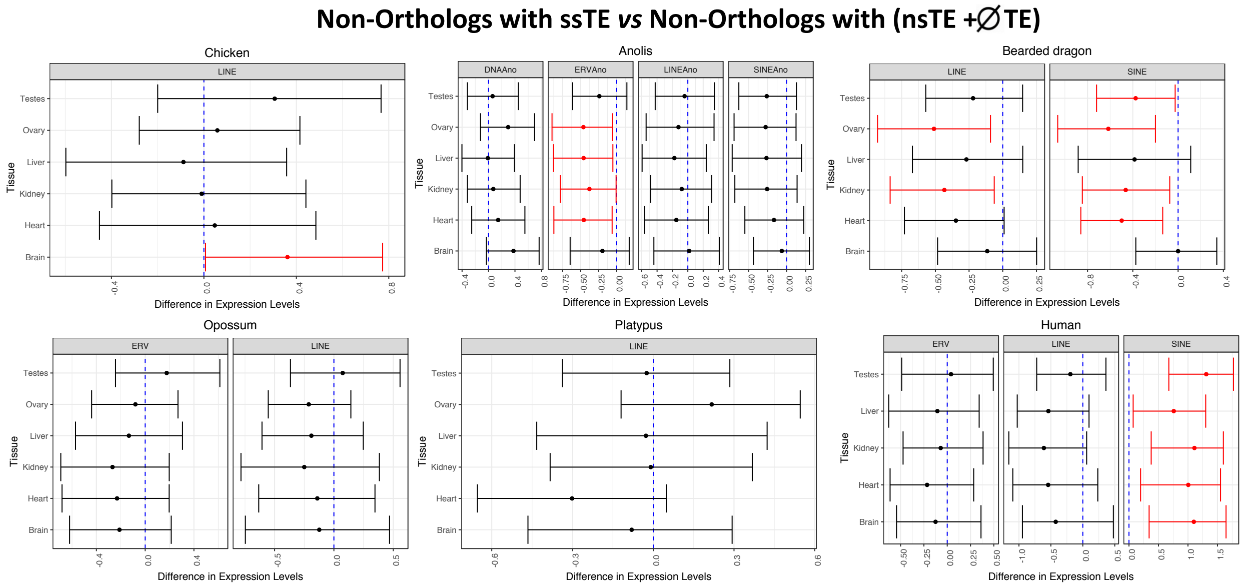

Figure S5: **Change in the level of non-ortholog gene expression as a function of species specific TE insertion.**
This figure shows the association between non-ortholog gene expression levels in six species (from left to right: anolis, chicken, human, opossum, platypus and bearded dragon (pogona)) with recent species-specific TE insertions (ssTE) (from left to right: LINE, SINE, ERV/LTR or DNA). Confidence Intervals for the difference in median log2(TPM) counts. Confidence Intervals were obtained using the weighted bootstrap and are 1-$\alpha$/m intervals, where $\alpha$=0.05 and m=nTissues*nElements as the total number of intervals presented. Red dots represent the median value from the bootstrap procedure, whilst the vertical line indicates zero. Intervals which do not contain zero are coloured red, and indicate a rejection of the null hypothesis, $H_0$: $\Delta\theta$=0, where $\theta$ represents the parameter of interest.

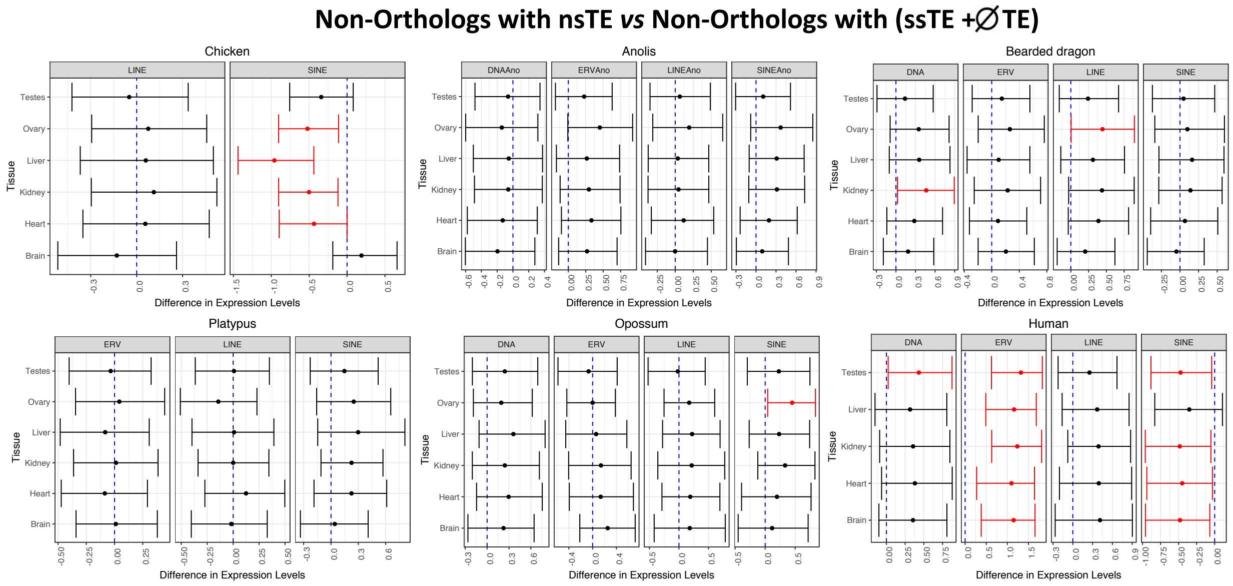

Figure S6: **Change in the level of non-ortholog gene expression as a function of non-species specific TE insertion.**
This figure shows the association between non-ortholog gene expression levels in six species (from left to right: anolis, chicken, human, opossum, platypus and bearded dragon (pogona)) with non-recent species-specific TE insertions (nsTE) (from left to right: LINE, SINE, ERV/LTR or DNA). Confidence Intervals for the difference in median log2(TPM) counts. Confidence Intervals were obtained using the weighted bootstrap and are 1-$\alpha$/m intervals, where $\alpha$=0.05 and m=nTissues*nElements as the total number of intervals presented. Red dots represent the median value from the bootstrap procedure, whilst the vertical line indicates zero. Intervals which do not contain zero are coloured red, and indicate a rejection of the null hypothesis, $H_0$: $\Delta\theta$=0, where $\theta$ represents the parameter of interest.

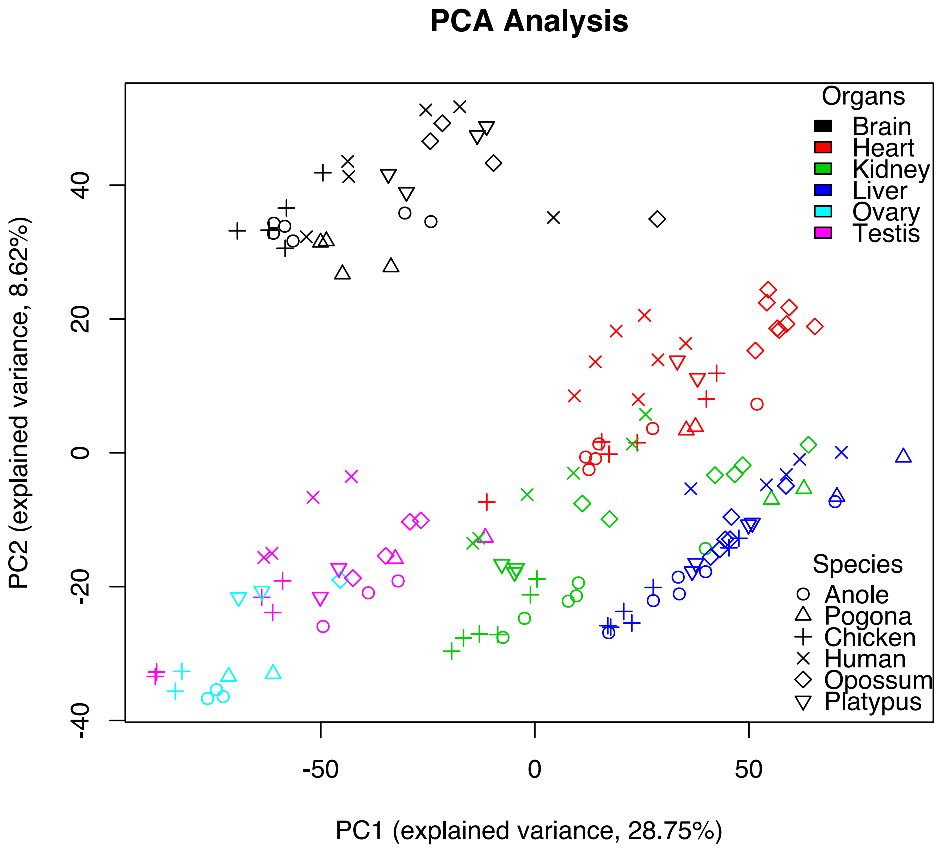

Figure S7: **Factorial map of the principal-component analysis of messenger RNA expression levels.**
This figure shows the PCA analysis of gene expression from six species (anole, bearded dragon (pogona), chicken, human, opossum and platypus) within six organs (brain, heart, kidney, liver, ovary and testis). Human samples did not include ovary. The proportion of the variance explained by the principal components is indicated in parentheses.

### Supplementary Tables

Table 1: **Gene expression dataset.** Show the systematic name, common name, Gender, Tissue, layout, source, study and instrument. Gene expression that were acquired through private collaboration (not publicly available) are marked as 'Private' in the Submitter column. The Following abbreviations are used for submitters:
IH2500 = Illumina HiSeq 2500
IH2000 = Illumina HiSeq 2000
IGA IIX = Illumina Genome Analyzer IIX

| Read accession(s) | Systematic Name | Common Name | Gender | Tissue | Layout | Source | Study | Instrument |
|---|---|---|---|---|---|---|---|---|
| SRR5412144 | *Anolis carolinensis* | Anole | Female | Brain | Single | NCBI | Marin | IH2500 |
| SRR5412145 | *Anolis carolinensis* | Anole | Female | Brain | Single | NCBI | Marin | IH2500 |
| SRR5412146 | *Anolis carolinensis* | Anole | Female | Brain | Single | NCBI | Marin | IH2500 |
| SRR5412147 | *Anolis carolinensis* | Anole | Male | Brain | Single | NCBI | Marin | IH2500 |
| SRR5412148 | *Anolis carolinensis* | Anole | Male | Brain | Single | NCBI | Marin | IH2500 |
| SRR5412149 | *Anolis carolinensis* | Anole | Male | Brain | Single | NCBI | Marin | IH2500 |
| SRR5412150 | *Anolis carolinensis* | Anole | Female | Heart | Single | NCBI | Marin | IH2500 |
| SRR5412151 | *Anolis carolinensis* | Anole | Female | Heart | Single | NCBI | Marin | IH2500 |
| SRR5412152 | *Anolis carolinensis* | Anole | Female | Heart | Single | NCBI | Marin | IH2500 |
| SRR5412153 | *Anolis carolinensis* | Anole | Male | Heart | Single | NCBI | Marin | IH2500 |
| SRR5412154 | *Anolis carolinensis* | Anole | Male | Heart | Single | NCBI | Marin | IH2500 |
| SRR5412155 | *Anolis carolinensis* | Anole | Male | Heart | Single | NCBI | Marin | IH2500 |
| SRR5412156 | *Anolis carolinensis* | Anole | Female | Kidney | Single | NCBI | Marin | IH2500 |
| SRR5412157 | *Anolis carolinensis* | Anole | Female | Kidney | Single | NCBI | Marin | IH2500 |
| SRR5412158 | *Anolis carolinensis* | Anole | Female | Kidney | Single | NCBI | Marin | IH2500 |
| SRR5412159 | *Anolis carolinensis* | Anole | Male | Kidney | Single | NCBI | Marin | IH2500 |
| SRR5412160 | *Anolis carolinensis* | Anole | Male | Kidney | Single | NCBI | Marin | IH2500 |
| SRR5412161 | *Anolis carolinensis* | Anole | Male | Kidney | Single | NCBI | Marin | IH2500 |
| SRR5412162 | *Anolis carolinensis* | Anole | Female | Liver | Single | NCBI | Marin | IH2500 |
| SRR5412163 | *Anolis carolinensis* | Anole | Female | Liver | Single | NCBI | Marin | IH2500 |
| SRR5412164 | *Anolis carolinensis* | Anole | Female | Liver | Single | NCBI | Marin | IH2500 |
| SRR5412165 | *Anolis carolinensis* | Anole | Male | Liver | Single | NCBI | Marin | IH2500 |
| SRR5412166 | *Anolis carolinensis* | Anole | Male | Liver | Single | NCBI | Marin | IH2500 |
| SRR5412167 | *Anolis carolinensis* | Anole | Male | Liver | Single | NCBI | Marin | IH2500 |
| SRR5412168 | *Anolis carolinensis* | Anole | Female | Ovary | Single | NCBI | Marin | IH2500 |
| SRR5412169 | *Anolis carolinensis* | Anole | Female | Ovary | Single | NCBI | Marin | IH2500 |
| SRR5412170 | *Anolis carolinensis* | Anole | Female | Ovary | Single | NCBI | Marin | IH2500 |
| SRR5412171 | *Anolis carolinensis* | Anole | Male | Testes | Single | NCBI | Marin | IH2500 |
| SRR5412172 | *Anolis carolinensis* | Anole | Male | Testes | Single | NCBI | Marin | IH2500 |
| SRR5412173 | *Anolis carolinensis* | Anole | Male | Testes | Single | NCBI | Marin | IH2500 |
| SRR5412242 | *Gallus gallus* | Chicken | Female | Brain | Single | NCBI | Marin | IH2500 |
| SRR5412243 | *Gallus gallus* | Chicken | Male | Brain | Single | NCBI | Marin | IH2500 |
| SRR5412244 | *Gallus gallus* | Chicken | Male | Brain | Single | NCBI | Marin | IH2500 |
| SRR306710 | *Gallus gallus* | Chicken | Female | Brain | Single | NCBI | Brawand | IGA IIX |
| SRR306711 | *Gallus gallus* | Chicken | Male | Brain | Single | NCBI | Brawand | IGA IIX |
| SRR5412245 | *Gallus gallus* | Chicken | Female | Heart | Single | NCBI | Marin | IH2500 |
| SRR5412246 | *Gallus gallus* | Chicken | Female | Heart | Single | NCBI | Marin | IH2500 |
| SRR5412247 | *Gallus gallus* | Chicken | Male | Heart | Single | NCBI | Marin | IH2500 |

| | | | | | | | | |
|---|---|---|---|---|---|---|---|---|
| SRR5412248 | *Gallus gallus* | Chicken | Male | Heart | Single | NCBI | Marin | IH2500 |
| SRR306714 | *Gallus gallus* | Chicken | Female | Heart | Single | NCBI | Brawand | IGA IIX |
| SRR306715 | *Gallus gallus* | Chicken | Male | Heart | Single | NCBI | Brawand | IGA IIX |
| SRR5412249 | *Gallus gallus* | Chicken | Female | Kidney | Single | NCBI | Marin | IH2500 |
| SRR5412250 | *Gallus gallus* | Chicken | Female | Kidney | Single | NCBI | Marin | IH2500 |
| SRR5412251 | *Gallus gallus* | Chicken | Male | Kidney | Single | NCBI | Marin | IH2500 |
| SRR5412252 | *Gallus gallus* | Chicken | Male | Kidney | Single | NCBI | Marin | IH2500 |
| SRR306716 | *Gallus gallus* | Chicken | Female | Kidney | Single | NCBI | Brawand | IGA IIX |
| SRR306717 | *Gallus gallus* | Chicken | Male | Kidney | Single | NCBI | Brawand | IGA IIX |
| SRR5412253 | *Gallus gallus* | Chicken | Female | Liver | Single | NCBI | Marin | IH2500 |
| SRR5412254 | *Gallus gallus* | Chicken | Female | Liver | Single | NCBI | Marin | IH2500 |
| SRR5412255 | *Gallus gallus* | Chicken | Male | Liver | Single | NCBI | Marin | IH2500 |
| SRR5412256 | *Gallus gallus* | Chicken | Male | Liver | Single | NCBI | Marin | IH2500 |
| SRR306718 | *Gallus gallus* | Chicken | Female | Liver | Single | NCBI | Brawand | IGA IIX |
| SRR306719 | *Gallus gallus* | Chicken | Male | Liver | Single | NCBI | Brawand | IGA IIX |
| SRR306720 | *Gallus gallus* | Chicken | Male | Liver | Single | NCBI | Brawand | IGA IIX |
| SRR5412257 | *Gallus gallus* | Chicken | Female | Ovary | Single | NCBI | Marin | IH2500 |
| SRR5412258 | *Gallus gallus* | Chicken | Female | Ovary | Single | NCBI | Marin | IH2500 |
| SRR5412259 | *Gallus gallus* | Chicken | Male | Testis | Single | NCBI | Marin | IH2500 |
| SRR5412260 | *Gallus gallus* | Chicken | Male | Testis | Single | NCBI | Marin | IH2500 |
| SRR306721 | *Gallus gallus* | Chicken | Male | Testis | Single | NCBI | Brawand | IGA IIX |
| SRR306722 | *Gallus gallus* | Chicken | Male | Testis | Single | NCBI | Brawand | IGA IIX |
| SRR306723 | *Gallus gallus* | Chicken | Male | Testis | Single | NCBI | Brawand | IGA IIX |
| ERR753525 | *Pogona Vitticeps* | Bearded dragon | Male | Brain | Paired | NCBI | Georges | IH2000 |
| ERR413064 | *Pogona Vitticeps* | Bearded dragon | Male | Brain | Paired | NCBI | Georges | IH2000 |
| ERR753526 | *Pogona Vitticeps* | Bearded dragon | Female | Brain | Paired | NCBI | Georges | IH2000 |
| ERR413071 | *Pogona Vitticeps* | Bearded dragon | Female | Brain | Paired | NCBI | Georges | IH2000 |
| ERR413072 | *Pogona Vitticeps* | Bearded dragon | Female | Heart | Paired | NCBI | Georges | IH2000 |
| ERR413065 | *Pogona Vitticeps* | Bearded dragon | Male | Heart | Paired | NCBI | Georges | IH2000 |
| ERR413073 | *Pogona Vitticeps* | Bearded dragon | Female | Kidney | Paired | NCBI | Georges | IH2000 |
| ERR413066 | *Pogona Vitticeps* | Bearded dragon | Male | Kidney | Paired | NCBI | Georges | IH2000 |
| ERR413074 | *Pogona Vitticeps* | Bearded dragon | Female | Liver | Paired | NCBI | Georges | IH2000 |
| ERR413067 | *Pogona Vitticeps* | Bearded dragon | Male | Liver | Paired | NCBI | Georges | IH2000 |
| ERR413070 | *Pogona Vitticeps* | Bearded dragon | Male | Testis | Paired | NCBI | Georges | IH2000 |
| ERR753529 | *Pogona Vitticeps* | Bearded dragon | Male | Testis | Paired | NCBI | Georges | IH2000 |
| ERR753530 | *Pogona Vitticeps* | Bearded dragon | Female | Ovary | Paired | NCBI | Georges | IH2000 |
| ERR413082 | *Pogona Vitticeps* | Bearded dragon | Female | Ovary | Paired | NCBI | Georges | IH2000 |
| SRR5412222 | *Ornithorhynchus anatinus* | Platypus | Female | Brain | Single | NCBI | Marin | IH2500 |
| SRR5412223 | *Ornithorhynchus anatinus* | Platypus | Female | Brain | Single | NCBI | Marin | IH2500 |
| SRR5412224 | *Ornithorhynchus anatinus* | Platypus | Male | Brain | Single | NCBI | Marin | IH2500 |
| SRR5412225 | *Ornithorhynchus anatinus* | Platypus | Male | Brain | Single | NCBI | Marin | IH2500 |
| SRR306724 | *Ornithorhynchus anatinus* | Platypus | Female | Brain | Single | NCBI | Brawand | IGA IIX |
| SRR306725 | *Ornithorhynchus anatinus* | Platypus | Female | Brain | Single | NCBI | Brawand | IGA IIX |
| SRR306726 | *Ornithorhynchus anatinus* | Platypus | Male | Brain | Single | NCBI | Brawand | IGA IIX |
| SRR306727 | *Ornithorhynchus anatinus* | Platypus | Male | Brain | Single | NCBI | Brawand | IGA IIX |
| SRR5412226 | *Ornithorhynchus anatinus* | Platypus | Female | Heart | Single | NCBI | Marin | IH2500 |

| | | | | | | | | |
|---|---|---|---|---|---|---|---|---|
| SRR5412227 | *Ornithorhynchus anatinus* | Platypus | Female | Heart | Single | NCBI | Marin | IH2500 |
| SRR5412228 | *Ornithorhynchus anatinus* | Platypus | Male | Heart | Single | NCBI | Marin | IH2500 |
| SRR5412229 | *Ornithorhynchus anatinus* | Platypus | Male | Heart | Single | NCBI | Marin | IH2500 |
| SRR306730 | *Ornithorhynchus anatinus* | Platypus | Female | Heart | Single | NCBI | Brawand | IGA IIX |
| SRR306731 | *Ornithorhynchus anatinus* | Platypus | Male | Heart | Single | NCBI | Brawand | IGA IIX |
| SRR5412230 | *Ornithorhynchus anatinus* | Platypus | Female | Kidney | Single | NCBI | Marin | IH2500 |
| SRR5412231 | *Ornithorhynchus anatinus* | Platypus | Female | Kidney | Single | NCBI | Marin | IH2500 |
| SRR5412232 | *Ornithorhynchus anatinus* | Platypus | Male | Kidney | Single | NCBI | Marin | IH2500 |
| SRR5412233 | *Ornithorhynchus anatinus* | Platypus | Male | Kidney | Single | NCBI | Marin | IH2500 |
| SRR306732 | *Ornithorhynchus anatinus* | Platypus | Female | Kidney | Single | NCBI | Brawand | IGA IIX |
| SRR306733 | *Ornithorhynchus anatinus* | Platypus | Male | Kidney | Single | NCBI | Brawand | IGA IIX |
| SRR306734 | *Ornithorhynchus anatinus* | Platypus | Male | Kidney | Single | NCBI | Brawand | IGA IIX |
| SRR5412234 | *Ornithorhynchus anatinus* | Platypus | Female | Liver | Single | NCBI | Marin | IH2500 |
| SRR5412235 | *Ornithorhynchus anatinus* | Platypus | Female | Liver | Single | NCBI | Marin | IH2500 |
| SRR5412236 | *Ornithorhynchus anatinus* | Platypus | Male | Liver | Single | NCBI | Marin | IH2500 |
| SRR5412237 | *Ornithorhynchus anatinus* | Platypus | Male | Liver | Single | NCBI | Marin | IH2500 |
| SRR306735 | *Ornithorhynchus anatinus* | Platypus | Female | Liver | Single | NCBI | Brawand | IGA IIX |
| SRR306736 | *Ornithorhynchus anatinus* | Platypus | Female | Liver | Single | NCBI | Brawand | IGA IIX |
| SRR306737 | *Ornithorhynchus anatinus* | Platypus | Male | Liver | Single | NCBI | Brawand | IGA IIX |
| SRR306738 | *Ornithorhynchus anatinus* | Platypus | Male | Liver | Single | NCBI | Brawand | IGA IIX |
| SRR5412238 | *Ornithorhynchus anatinus* | Platypus | Female | Ovary | Single | NCBI | Marin | IH2500 |
| SRR5412239 | *Ornithorhynchus anatinus* | Platypus | Female | Ovary | Single | NCBI | Marin | IH2500 |
| SRR5412240 | *Ornithorhynchus anatinus* | Platypus | Male | Testis | Single | NCBI | Marin | IH2500 |
| SRR5412241 | *Ornithorhynchus anatinus* | Platypus | Male | Testis | Single | NCBI | Marin | IH2500 |
| SRR306739 | *Ornithorhynchus anatinus* | Platypus | Male | Testis | Single | NCBI | Brawand | IGA IIX |
| SRR306741 | *Ornithorhynchus anatinus* | Platypus | Male | Testis | Single | NCBI | Brawand | IGA IIX |
| SRR5412205 | *Monodelphis domestica* | Opossum | Female | Brain | Single | NCBI | Marin | IH2500 |
| SRR5412206 | *Monodelphis domestica* | Opossum | Male | Brain | Single | NCBI | Marin | IH2500 |
| SRR306742 | *Monodelphis domestica* | Opossum | Female | Brain | Single | NCBI | Brawand | IGA IIX |
| SRR306743 | *Monodelphis domestica* | Opossum | Female | Brain | Single | NCBI | Brawand | IGA IIX |
| SRR306744 | *Monodelphis domestica* | Opossum | Male | Brain | Single | NCBI | Brawand | IGA IIX |
| SRR5412207 | *Monodelphis domestica* | Opossum | Female | Heart | Single | NCBI | Marin | IH2500 |
| SRR5412208 | *Monodelphis domestica* | Opossum | Female | Heart | Single | NCBI | Marin | IH2500 |
| SRR5412209 | *Monodelphis domestica* | Opossum | Male | Heart | Single | NCBI | Marin | IH2500 |
| SRR5412210 | *Monodelphis domestica* | Opossum | Male | Heart | Single | NCBI | Marin | IH2500 |
| SRR306747 | *Monodelphis domestica* | Opossum | Female | Heart | Single | NCBI | Brawand | IGA IIX |
| SRR306748 | *Monodelphis domestica* | Opossum | Female | Heart | Single | NCBI | Brawand | IGA IIX |
| SRR306749 | *Monodelphis domestica* | Opossum | Male | Heart | Single | NCBI | Brawand | IGA IIX |
| SRR306750 | *Monodelphis domestica* | Opossum | Male | Heart | Single | NCBI | Brawand | IGA IIX |
| SRR5412211 | *Monodelphis domestica* | Opossum | Female | Kidney | Single | NCBI | Marin | IH2500 |
| SRR5412212 | *Monodelphis domestica* | Opossum | Female | Kidney | Single | NCBI | Marin | IH2500 |
| SRR5412213 | *Monodelphis domestica* | Opossum | Male | Kidney | Single | NCBI | Marin | IH2500 |
| SRR5412214 | *Monodelphis domestica* | Opossum | Male | Kidney | Single | NCBI | Marin | IH2500 |
| SRR306751 | *Monodelphis domestica* | Opossum | Female | Kidney | Single | NCBI | Brawand | IGA IIX |
| SRR306752 | *Monodelphis domestica* | Opossum | Male | Kidney | Single | NCBI | Brawand | IGA IIX |
| SRR5412215 | *Monodelphis domestica* | Opossum | Female | Liver | Single | NCBI | Marin | IH2500 |
| SRR5412216 | *Monodelphis domestica* | Opossum | Female | Liver | Single | NCBI | Marin | IH2500 |
| SRR5412217 | *Monodelphis domestica* | Opossum | Male | Liver | Single | NCBI | Marin | IH2500 |
| SRR5412218 | *Monodelphis domestica* | Opossum | Male | Liver | Single | NCBI | Marin | IH2500 |
| SRR306753 | *Monodelphis domestica* | Opossum | Female | Liver | Single | NCBI | Brawand | IGA IIX |

| | | | | | | | | |
|---|---|---|---|---|---|---|---|---|
| SRR306754 | *Monodelphis domestica* | Opossum | Male | Liver | Single | NCBI | Brawand | IGA IIX |
| SRR5412219 | *Monodelphis domestica* | Opossum | Female | Ovary | Single | NCBI | Marin | IH2500 |
| SRR5412220 | *Monodelphis domestica* | Opossum | Male | Testis | Single | NCBI | Marin | IH2500 |
| SRR5412221 | *Monodelphis domestica* | Opossum | Male | Testis | Single | NCBI | Marin | IH2500 |
| SRR306755 | *Monodelphis domestica* | Opossum | Male | Testis | Single | NCBI | Brawand | IGA IIX |
| SRR306756 | *Monodelphis domestica* | Opossum | Male | Testis | Single | NCBI | Brawand | IGA IIX |
| SRR5412174 | *Homo sapiens* | Human | Female | Brain | Single | NCBI | Marin | IH2500 |
| SRR5412175 | *Homo sapiens* | Human | Male | Brain | Single | NCBI | Marin | IH2500 |
| SRR306838 | *Homo sapiens* | Human | Female | Brain | Single | NCBI | Brawand | IGA IIX |
| SRR306839 | *Homo sapiens* | Human | Male | Brain | Single | NCBI | Brawand | IGA IIX |
| SRR306841 | *Homo sapiens* | Human | Male | Brain | Single | NCBI | Brawand | IGA IIX |
| SRR306843 | *Homo sapiens* | Human | Male | Brain | Single | NCBI | Brawand | IGA IIX |
| SRR5412176 | *Homo sapiens* | Human | Female | Heart | Paired | NCBI | Marin | IH2500 |
| SRR5412177 | *Homo sapiens* | Human | Male | Heart | Single | NCBI | Marin | IH2500 |
| SRR5412178 | *Homo sapiens* | Human | Male | Heart | Paired | NCBI | Marin | IH2500 |
| SRR306847 | *Homo sapiens* | Human | Female | Heart | Single | NCBI | Brawand | IGA IIX |
| SRR306848 | *Homo sapiens* | Human | Male | Heart | Single | NCBI | Brawand | IGA IIX |
| SRR306849 | *Homo sapiens* | Human | Male | Heart | Single | NCBI | Brawand | IGA IIX |
| SRR306850 | *Homo sapiens* | Human | Male | Heart | Single | NCBI | Brawand | IGA IIX |
| SRR5412179 | *Homo sapiens* | Human | Female | Kidney | Single | NCBI | Marin | IH2500 |
| SRR5412180 | *Homo sapiens* | Human | Male | Kidney | Single | NCBI | Marin | IH2500 |
| SRR5412181 | *Homo sapiens* | Human | Male | Kidney | Single | NCBI | Marin | IH2500 |
| SRR306851 | *Homo sapiens* | Human | Female | Kidney | Single | NCBI | Brawand | IGA IIX |
| SRR306852 | *Homo sapiens* | Human | Male | Kidney | Single | NCBI | Brawand | IGA IIX |
| SRR306853 | *Homo sapiens* | Human | Male | Kidney | Single | NCBI | Brawand | IGA IIX |
| SRR5412182 | *Homo sapiens* | Human | Female | Liver | Single | NCBI | Marin | IH2500 |
| SRR5412183 | *Homo sapiens* | Human | Male | Liver | Single | NCBI | Marin | IH2500 |
| SRR306854 | *Homo sapiens* | Human | Male | Liver | Single | NCBI | Brawand | IGA IIX |
| SRR306855 | *Homo sapiens* | Human | Male | Liver | Single | NCBI | Brawand | IGA IIX |
| SRR306856 | *Homo sapiens* | Human | Male | Liver | Single | NCBI | Brawand | IGA IIX |
| SRR5412184 | *Homo sapiens* | Human | Male | Testis | Single | NCBI | Marin | IH2500 |
| SRR5412185 | *Homo sapiens* | Human | Male | Testis | Single | NCBI | Marin | IH2500 |
| SRR306857 | *Homo sapiens* | Human | Male | Testes | Single | NCBI | Brawand | IGA IIX |
| SRR306858 | *Homo sapiens* | Human | Male | Testes | Single | NCBI | Brawand | IGA IIX |

Table 2: **Assembly dataset**. Shows the systematic name, common name, genome version, source and submitter for all the genomes tested with our *ab initio* method.
The Following abbreviations are used for submitters:
Genome Sequencing Platform, The Genome Assembly Team = GAT;
Genome Reference Consortium = GRC;
International Chicken Genome Consortium = ICGS;
Washington University = WashU.

| No | Systematic Name | Common Name | RefSeq Assembly Accession | Source | Submitter |
|---|---|---|---|---|---|
| 1 | *Homo sapiens* | Human | GCF_000001405.25 | NCBI | GRC |
| 2 | *Pogona Vitticeps* | Bearded Dragon | GCF_900067755.1 | NCBI | BRAEMBL |
| 3 | *Anolis Carollinensis* | Anolis lizard | GCF_000090745.1 | NCBI | Broad |
| 4 | *Gallus gallus* | Chicken | GCF_000002315.3 | NCBI | ICGS |
| 5 | *Monodelphis domestica* | Opossum | GCF_000002295.2 | NCBI | GAT |
| 6 | *Ornithorhynchus anatinus* | Platypus | GCF_000002275.2 | NCBI | WashU |

Table 3: **Comparsion of orthologs with ssTE *vs* orthologs with nsTE and ∅ TE**. Shows the number of sample genes used in the bootstrap approach. Test sample is ortholog genes containing recent species specific TE (ssTE), reference sample is ortholog genes with no ssTE.

| | **Chicken** | | **Anolis** | | **Bearded dragon** | | **Platypus** | | **Opossum** | | **Human** | |
|---|---|---|---|---|---|---|---|---|---|---|---|---|
| | Test | Reference | Test | Reference | Test | Reference | Test | Reference | Test | Reference | Test | Reference |
| LINE | 1,580 | 5,015 | 4,135 | 2,640 | 3,613 | 2,982 | 1,854 | 4,741 | 3,274 | 3,321 | 2,048 | 4,547 |
| SINE | 0 | NA | 1,566 | 5,029 | 5,660 | 935 | 513 | NA | 317 | NA | 3,388 | 3,207 |
| ERV | 143 | NA | 2,340 | 4,255 | 104 | NA | 16 | NA | 3,064 | 3,531 | 994 | 5,601 |
| DNA | 5 | NA | 3,436 | 3,159 | 496 | NA | 6 | NA | 236 | NA | 45 | NA |

Table 4: **Comparsion of orthologs with nsTE *vs* orthologs with ssTE and ∅ TE**. Shows the number of sample genes used in the bootstrap approach. Test sample is ortholog genes containing non-recent species specific TE (nsTE), reference sample is ortholog genes with no nsTE.

| | **Chicken** | | **Anolis** | | **Bearded dragon** | | **Platypus** | | **Opossum** | | **Human** | |
|---|---|---|---|---|---|---|---|---|---|---|---|---|
| | Test | Reference | Test | Reference | Test | Reference | Test | Reference | Test | Reference | Test | Reference |
| LINE | 4,320 | 2,275 | 2,221 | 4,374 | 2,805 | 3,790 | 4,516 | 2,079 | 3,013 | 3,582 | 4,340 | 2,255 |
| SINE | 1,174 | 5,421 | 4,369 | 2,226 | 4,667 | 1,928 | 5,830 | 765 | 6,076 | NA | 3,070 | 3,525 |
| ERV | 5,797 | NA | 3,652 | 2,943 | 6,106 | NA | 5,374 | NA | 2,871 | 3,724 | 5,470 | 1,125 |
| DNA | 5,894 | NA | 3,066 | 3,529 | 5,931 | NA | 5,525 | NA | 5,819 | NA | 6,455 | NA |

Table 5: **Comparsion of non-orthologs with ssTE *vs* non-orthologs with nsTE and ∅ TE**. Shows the number of sample genes used in bootstrap approach. Test sample is non-ortholog genes containing recent species specific TE (ssTE), reference sample is non-ortholog genes with no ssTE.

| | **Chicken** | | **Anolis** | | **Bearded dragon** | | **Platypus** | | **Opossum** | | **Human** | |
|---|---|---|---|---|---|---|---|---|---|---|---|---|
| | Test | Reference | Test | Reference | Test | Reference | Test | Reference | Test | Reference | Test | Reference |
| LINE | 1,488 | 9,025 | 8,337 | 10,988 | 5,671 | 9,728 | 2,677 | 16,844 | 5,025 | 12,279 | 6,065 | 45,076 |
| SINE | 0 | NA | 2,670 | 16,655 | 1,203 | 14,196 | 553 | NA | 413 | NA | 8,603 | 42,538 |
| ERV | 211 | NA | 4,528 | 14,797 | 142 | NA | 24 | NA | 4,439 | 12,865 | 3,401 | 47,740 |
| DNA | 11 | NA | 5,593 | 13,372 | 560 | NA | 5 | NA | 344 | NA | 220 | NA |

Table 6: **Comparsion of non-orthologs with nsTE *vs* non-orthologs with ssTE and ∅ TE**. Shows the number of sample genes used in the bootstrap approach. Test sample is non-ortholog genes containing non-recent species specific TE (nsTE), reference sample is non-ortholog genes with no nsTE.

| | **Chicken** | | **Anolis** | | **Bearded dragon** | | **Platypus** | | **Opossum** | | **Human** | |
|---|---|---|---|---|---|---|---|---|---|---|---|---|
| | Test | Reference | Test | Reference | Test | Reference | Test | Reference | Test | Reference | Test | Reference |
| LINE | 6,402 | 4,111 | 8,186 | 11,139 | 8,428 | 6,971 | 14,113 | 5,408 | 9,423 | 7,881 | 32,875 | 18,266 |
| SINE | 1,191 | 9,322 | 11,472 | 7,853 | 9,147 | 6,252 | 16,175 | 3,346 | 13,241 | 4,063 | 32,869 | 18,272 |
| ERV | 7,453 | 3,060 | 8,690 | 10,635 | 12,671 | 2,728 | 9,778 | 9,743 | 7,360 | 9,944 | 34,177 | 18,202 |
| DNA | 7,397 | 3,116 | 11,320 | 8,005 | 13,190 | 2,209 | 9,878 | NA | 10,232 | 7,072 | 32,939 | 16,964 |

Table 7: **Difference in the gene expression of orthologs/non-orthologs with a TE insertion.** Shows the species, TE element, Tissue, gene expression comparison sets , lowest gene expression level, median gene expression level, highest gene expression level, bonferroni-adjusted 95% CI lowest gene expression, bonferroni-adjusted 95% CI highest expression level and the significance indicator. TPM counts were log2 transformed.

| Species | Element | Tissue | Data | lwr | med | upr | lwr95 | upr95 | Sig |
|---|---|---|---|---|---|---|---|---|---|
| Platypus | LINE | Heart | ssTE-ortholog | -0.859 | -0.443 | -0.045 | -0.769 | -0.131 | T |
| Pogona | LINE | Heart | ssTE-ortholog | -0.704 | -0.387 | -0.064 | -0.626 | -0.147 | T |
| Pogona | LINE | Kidney | ssTE-ortholog | -0.815 | -0.485 | -0.160 | -0.732 | -0.242 | T |
| Pogona | LINE | Liver | ssTE-ortholog | -0.864 | -0.525 | -0.188 | -0.771 | -0.276 | T |
| Pogona | LINE | Ovary | ssTE-ortholog | -0.975 | -0.629 | -0.278 | -0.883 | -0.373 | T |
| Pogona | LINE | Testes | ssTE-ortholog | -0.677 | -0.369 | -0.041 | -0.599 | -0.129 | T |
| Pogona | SINE | Heart | ssTE-ortholog | -0.946 | -0.584 | -0.218 | -0.851 | -0.322 | T |
| Pogona | SINE | Kidney | ssTE-ortholog | -1.035 | -0.672 | -0.325 | -0.933 | -0.406 | T |
| Pogona | SINE | Liver | ssTE-ortholog | -0.965 | -0.566 | -0.194 | -0.851 | -0.283 | T |
| Pogona | SINE | Ovary | ssTE-ortholog | -1.022 | -0.592 | -0.212 | -0.901 | -0.300 | T |
| Pogona | SINE | Testes | ssTE-ortholog | -0.939 | -0.601 | -0.259 | -0.854 | -0.351 | T |
| Chicken | LINE | Brain | ssTE-ortholog | 0.013 | 0.323 | 0.667 | 0.089 | 0.579 | T |
| Anolis | LINE | Heart | ssTE-ortholog | -0.792 | -0.444 | -0.112 | -0.701 | -0.197 | T |
| Anolis | LINE | Kidney | ssTE-ortholog | -0.830 | -0.459 | -0.128 | -0.724 | -0.212 | T |
| Anolis | LINE | Liver | ssTE-ortholog | -0.832 | -0.481 | -0.120 | -0.743 | -0.211 | T |
| Anolis | LINE | Ovary | ssTE-ortholog | -0.730 | -0.402 | -0.087 | -0.647 | -0.168 | T |
| Anolis | SINE | Heart | ssTE-ortholog | -0.847 | -0.509 | -0.191 | -0.758 | -0.266 | T |
| Anolis | SINE | Kidney | ssTE-ortholog | -0.824 | -0.506 | -0.185 | -0.745 | -0.267 | T |
| Anolis | SINE | Ovary | ssTE-ortholog | -0.838 | -0.493 | -0.170 | -0.751 | -0.252 | T |
| Anolis | SINE | Testes | ssTE-ortholog | -0.762 | -0.459 | -0.154 | -0.684 | -0.232 | T |
| Anolis | ERV | Heart | ssTE-ortholog | -0.760 | -0.426 | -0.104 | -0.669 | -0.187 | T |
| Anolis | ERV | Kidney | ssTE-ortholog | -0.968 | -0.653 | -0.348 | -0.886 | -0.425 | T |
| Anolis | ERV | Liver | ssTE-ortholog | -1.011 | -0.680 | -0.347 | -0.924 | -0.433 | T |
| Anolis | ERV | Ovary | ssTE-ortholog | -0.709 | -0.391 | -0.066 | -0.628 | -0.151 | T |
| Human | LINE | Kidney | ssTE-ortholog | -1.179 | -0.746 | -0.041 | -1.079 | -0.126 | T |
| Human | LINE | Liver | ssTE-ortholog | -1.283 | -0.738 | -0.116 | -1.179 | -0.216 | T |
| Human | LINE | Testes | ssTE-ortholog | -0.917 | -0.562 | -0.221 | -0.840 | -0.296 | T |
| Human | SINE | Brain | ssTE-ortholog | 0.119 | 0.429 | 0.750 | 0.196 | 0.671 | T |
| Human | SINE | Testes | ssTE-ortholog | 0.274 | 0.619 | 0.933 | 0.357 | 0.858 | T |
| Pogona | LINE | Kidney | ssTE-nonOrtholog | -0.836 | -0.433 | -0.063 | -0.695 | -0.172 | T |
| Pogona | LINE | Ovary | ssTE-nonOrtholog | -0.929 | -0.510 | -0.091 | -0.797 | -0.226 | T |
| Pogona | SINE | Heart | ssTE-nonOrtholog | -0.859 | -0.498 | -0.136 | -0.748 | -0.250 | T |
| Pogona | SINE | Kidney | ssTE-nonOrtholog | -0.846 | -0.463 | -0.074 | -0.725 | -0.199 | T |
| Pogona | SINE | Ovary | ssTE-nonOrtholog | -1.063 | -0.616 | -0.200 | -0.933 | -0.334 | T |
| Pogona | SINE | Testes | ssTE-nonOrtholog | -0.720 | -0.375 | -0.026 | -0.619 | -0.136 | T |
| Chicken | LINE | Brain | ssTE-nonOrtholog | 0.008 | 0.388 | 0.832 | 0.125 | 0.694 | T |
| Anolis | ERV | Heart | ssTE-nonOrtholog | -0.855 | -0.456 | -0.076 | -0.730 | -0.191 | T |
| Anolis | ERV | Kidney | ssTE-nonOrtholog | -0.764 | -0.378 | -0.020 | -0.640 | -0.127 | T |
| Anolis | ERV | Liver | ssTE-nonOrtholog | -0.865 | -0.458 | -0.062 | -0.737 | -0.191 | T |
| Anolis | ERV | Ovary | ssTE-nonOrtholog | -0.876 | -0.461 | -0.073 | -0.743 | -0.194 | T |
| Human | SINE | Brain | ssTE-nonOrtholog | 0.350 | 1.099 | 1.629 | 0.470 | 1.478 | T |
| Human | SINE | Heart | ssTE-nonOrtholog | 0.206 | 1.005 | 1.538 | 0.315 | 1.375 | T |
| Human | SINE | Kidney | ssTE-nonOrtholog | 0.386 | 1.109 | 1.590 | 0.511 | 1.451 | T |
| Human | SINE | Liver | ssTE-nonOrtholog | 0.085 | 0.759 | 1.292 | 0.218 | 1.147 | T |
| Human | SINE | Testes | ssTE-nonOrtholog | 0.685 | 1.310 | 1.765 | 0.815 | 1.621 | T |
| Platypus | SINE | Liver | nsTE-ortholog | 0.089 | 0.622 | 1.168 | 0.251 | 1.001 | T |
| Platypus | SINE | Ovary | nsTE-ortholog | 0.027 | 0.473 | 0.924 | 0.162 | 0.780 | T |
| Platypus | SINE | Testes | nsTE-ortholog | 0.114 | 0.567 | 1.031 | 0.250 | 0.890 | T |
| Pogona | LINE | Heart | nsTE-ortholog | 0.009 | 0.391 | 0.786 | 0.134 | 0.649 | T |

| | | | | | | | | | |
|---|---|---|---|---|---|---|---|---|---|
| Pogona | LINE | Kidney | nsTE-ortholog | 0.050 | 0.428 | 0.795 | 0.172 | 0.687 | T |
| Pogona | LINE | Liver | nsTE-ortholog | 0.125 | 0.523 | 0.916 | 0.250 | 0.790 | T |
| Pogona | LINE | Ovary | nsTE-ortholog | 0.186 | 0.577 | 0.972 | 0.300 | 0.840 | T |
| Pogona | LINE | Testes | nsTE-ortholog | 0.007 | 0.368 | 0.733 | 0.125 | 0.610 | T |
| Chicken | SINE | Heart | nsTE-ortholog | -1.039 | -0.624 | -0.250 | -0.905 | -0.365 | T |
| Chicken | SINE | Kidney | nsTE-ortholog | -1.134 | -0.743 | -0.356 | -1.012 | -0.472 | T |
| Chicken | SINE | Liver | nsTE-ortholog | -1.424 | -0.935 | -0.451 | -1.272 | -0.583 | T |
| Chicken | SINE | Ovary | nsTE-ortholog | -1.051 | -0.685 | -0.340 | -0.935 | -0.447 | T |
| Chicken | SINE | Testes | nsTE-ortholog | -1.025 | -0.715 | -0.412 | -0.925 | -0.508 | T |
| Anolis | LINE | Heart | nsTE-ortholog | 0.149 | 0.514 | 0.895 | 0.260 | 0.777 | T |
| Anolis | LINE | Kidney | nsTE-ortholog | 0.212 | 0.567 | 0.974 | 0.320 | 0.844 | T |
| Anolis | LINE | Liver | nsTE-ortholog | 0.175 | 0.578 | 0.978 | 0.303 | 0.852 | T |
| Anolis | LINE | Ovary | nsTE-ortholog | 0.170 | 0.534 | 0.919 | 0.282 | 0.795 | T |
| Anolis | SINE | Heart | nsTE-ortholog | 0.035 | 0.393 | 0.758 | 0.148 | 0.643 | T |
| Anolis | SINE | Ovary | nsTE-ortholog | 0.109 | 0.474 | 0.836 | 0.222 | 0.726 | T |
| Anolis | SINE | Testes | nsTE-ortholog | 0.084 | 0.431 | 0.773 | 0.196 | 0.665 | T |
| Human | LINE | Liver | nsTE-ortholog | 0.033 | 0.584 | 1.159 | 0.155 | 1.002 | T |
| Human | LINE | Testes | nsTE-ortholog | 0.102 | 0.474 | 0.844 | 0.209 | 0.732 | T |
| Human | SINE | Testes | nsTE-ortholog | -0.776 | -0.411 | -0.063 | -0.664 | -0.169 | T |
| Pogona | LINE | Ovary | nsTE-nonOrtholog | 0.012 | 0.446 | 0.879 | 0.155 | 0.743 | T |
| Pogona | DNA | Kidney | nsTE-nonOrtholog | 0.023 | 0.464 | 0.892 | 0.186 | 0.751 | T |
| Chicken | SINE | Kidney | nsTE-nonOrtholog | -0.915 | -0.502 | -0.102 | -0.775 | -0.234 | T |
| Chicken | SINE | Liver | nsTE-nonOrtholog | -1.459 | -0.960 | -0.421 | -1.292 | -0.585 | T |
| Chicken | SINE | Ovary | nsTE-nonOrtholog | -0.913 | -0.522 | -0.086 | -0.781 | -0.255 | T |
| Opossum | SINE | Ovary | nsTE-nonOrtholog | 0.053 | 0.449 | 0.830 | 0.178 | 0.709 | T |
| Human | SINE | Brain | nsTE-nonOrtholog | -0.955 | -0.481 | -0.071 | -0.803 | -0.198 | T |
| Human | SINE | Heart | nsTE-nonOrtholog | -0.937 | -0.451 | -0.033 | -0.772 | -0.156 | T |
| Human | SINE | Kidney | nsTE-nonOrtholog | -0.955 | -0.485 | -0.054 | -0.804 | -0.191 | T |
| Human | SINE | Testes | nsTE-nonOrtholog | -0.879 | -0.473 | -0.044 | -0.740 | -0.202 | T |
| Human | ERV | Brain | nsTE-nonOrtholog | 0.382 | 1.149 | 1.651 | 0.521 | 1.482 | T |
| Human | ERV | Heart | nsTE-nonOrtholog | 0.278 | 1.097 | 1.638 | 0.437 | 1.469 | T |
| Human | ERV | Kidney | nsTE-nonOrtholog | 0.631 | 1.235 | 1.806 | 0.787 | 1.611 | T |
| Human | ERV | Liver | nsTE-nonOrtholog | 0.493 | 1.159 | 1.683 | 0.636 | 1.521 | T |
| Human | ERV | Testes | nsTE-nonOrtholog | 0.624 | 1.321 | 1.815 | 0.765 | 1.656 | T |
| Human | DNA | Testes | nsTE-nonOrtholog | 0.020 | 0.433 | 0.891 | 0.162 | 0.717 | T |
